# Designing Distributed Cell Classifier Circuits using a Genetic Algorithm

**DOI:** 10.1101/652339

**Authors:** Melania Nowicka, Heike Siebert

## Abstract

Cell classifiers are decision-making synthetic circuits that allow in vivo cell-type classification. Their design is based on finding a relationship between differential expression of miRNAs and the cell condition. Such biological devices have shown potential to become a valuable tool in cancer treatment as a new type-specific cell targeting approach. So far, only single-circuit classifiers were designed in this context. However, reliable designs come with high complexity, making them difficult to assemble in the lab. Here, we apply so-called Distributed Classifiers (DC) consisting of simple single circuits, that decide collectively according to a threshold function. Such architecture potentially simplifies the assembly process and provides design flexibility. Here, we present a genetic algorithm that allows the design and optimization of DCs. Breast cancer case studies show that DCs perform with high accuracy on real-world data. Optimized classifiers capture biologically relevant miRNAs that are cancer-type specific. The comparison to a single-circuit classifier design approach shows that DCs perform with significantly higher accuracy than individual circuits. The algorithm is implemented as an open source tool.

## 1 Introduction

Synthetic biology has shown its immense potential in recent years in a wide array of applications. This is particularly true for the medical field, where synthetic biological systems are developed for versatile employment from diagnostics to treatment [29, 25]. Research in design and construction of cell classifier circuits touches on both these areas. Cell classifiers are molecular constructs capable of sensing certain markers in the environment, processing the input and reacting with a signal-specific output. A prime example for this are miRNA-based classifiers that distinguish cell states, e.g., as healthy or diseased, based on their miRNA expression profiles applying boolean logic (Fig. 1A) [27, 16]. These circuits can be delivered to cells on plasmids or viral vectors and trigger the production of a desired output, e.g., a toxic compound causing cell apoptosis in diseased cells (Fig. 1B).

**Fig. 1.**
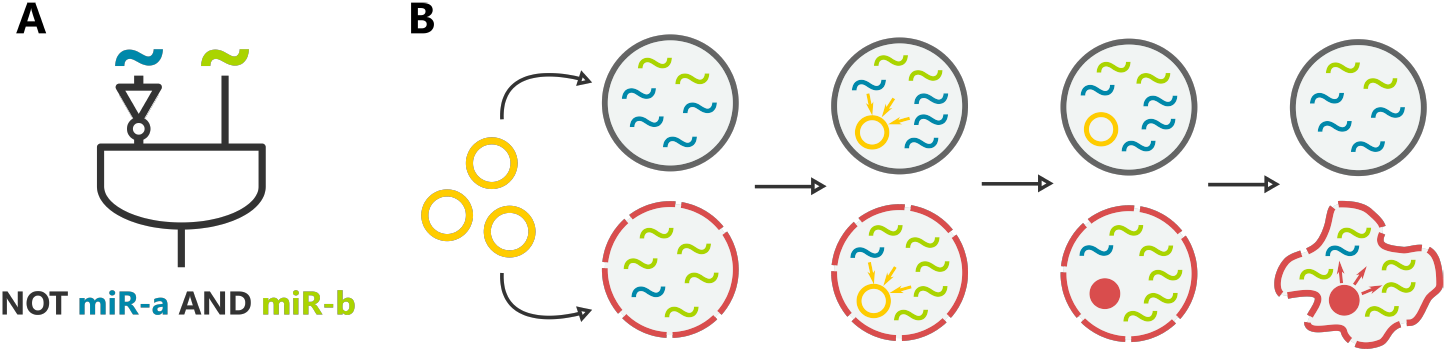
(A) An exemplary boolean design of a two miRNA-input cell classifier. (B) A schema showing two types of cells, healthy and diseased (dashed line). The classifiers are delivered to the cells, sense the internal input levels and respond with respect to a given cell condition.

A variety of different approaches to designing synthetic circuits is available [18, 13, 26]. However, to confront many application-derived limitations, circuit designs must be often tailored to rigorous specifications. Since cell classifiers must be feasible to implement in the lab, many constraints are posed on the building blocks of these circuits that need to be encoded in the design problem. So far, two different methods for single-circuit classifiers were described [16, 1]. Mohammadi et al. [16] proposed a heuristic approach that allows to optimize a classifier’s topology using a mechanistic model of the circuit and a predefined set of biochemical parameters. Another approach was presented by Becker et al. [1]. The authors propose a method for finding globally optimal classifiers represented by boolean functions based on binarized miRNA expression data. To search through the entire space of solutions in a short time frame the authors apply logic solvers. Becker et al. compare their results to the previously mentioned state-of-the-art method demonstrating significant improvement in binary classification of presented classifiers [1].

While this research shows that theoretically single-circuit classifiers can perform such classification tasks [16, 1], there is a number of challenges for the approach in application. Depending on the heterogeneity of the data, to obtain a clear-cut classification often a circuit of high complexity is needed. Generally, the cost both in time and money for classifier circuit construction in the lab goes up the larger and more complex the circuit architecture gets, quickly becoming not feasible at all [16]. A further problem is the robustness needed for reliable performance when faced with uncertainty and noise in signals and wide ranging possibilities for perturbations of the classifier functionality in natural environments. To address these issues the principles of distributed classification, as inherent in many natural systems such as the immune system and shown to be an effective strategy, e.g., in machine learning, can be exploited [23, 20]. Here, the idea is to design a set of different classifier circuits, also called distributed classifier, that perform classification in an integrated manner. Such a set can consist of rather simple classifiers that still perform better and more robustly than a complex single circuit classifier, since the individual classification results are aggregated which compensates for individual mistakes. A theoretical design of such a distributed classifier based on synthetic gene circuits was presented by Didovyk et al. [3]. The classifier is optimized by training a starting population of simple circuits on the available data similarly to machine learning algorithms, i.e., by presenting learning examples and successively removing low-performance circuits. While this work considers only a quite specific scenario and the approach has some drawbacks such as strong dependence on the quality of the starting population, it highlights the potential of the underlying idea of using distributed classifiers.

Here, we adapt the distributed classifier approach proposed by Didovyk et al. [3] to the problem of cell classifier design. We define a *Distributed Classifier* (DC) as a set of single-circuit classifiers that decide collectively based on a threshold function. Biologically, the threshold may correspond to a certain concentration of the drug that allows to treat the cells or fluorescent marker allowing to classify the cell type [3, 16, 15]. According to Mohammadi et al. [16] such threshold manipulation may be achieved by changing the biochemical parameters of a circuit model. Due to the high complexity of the problem, we apply a heuristic approach to design and optimize DCs, namely, a genetic algorithm (GA). GAs are evolution-inspired metaheuristics that allow to optimize populations of individuals [14]. Such evolutionary approaches were successfully applied to various biological questions [12], e.g., design of synthetic networks and, in particular, design of single-circuit classifiers [26, 16]. Due to the high flexibility of GAs in terms of design and parameters, the algorithm may be efficiently adapted to the distributed classifier problem.

In this article, we illustrate the potential of distributed classifiers in application, in particular, in cancer cell classification. The following section contains preliminaries including the definition of a single-circuit and distributed classifier. Section 3 describes the architecture of the proposed genetic algorithm for the design and optimization of DCs. In Section 4 we present case studies performed on real-world breast cancer data and compare the results with a single-circuit design method proposed by Becker et al. [1]. Finally, we discuss the distributed classifier performance and comment on potential future work.

## 2 Preliminaries

In this section we describe the data we employ to designing classifiers, introduce single-circuit and distributed classifiers and propose binary classification measures that allow to evaluate their performance.

### 2.1 miRNA Expression Data

The proposed method is a boolean approach and utilizes binarized and annotated data. While our focus is on miRNA expression profiles, the approach can naturally be applied to any data set of the format introduced below.

In cancer research, differentially expressed miRNAs provide a valuable source of information about tumor development, progression and response to a therapy [10, 9]. Thus, dysregulated miRNAs have been considered as potential biomarkers for cancer diagnosis and treatment. One of the approaches allowing to distinguish up- and down-regulated miRNAs is discretization of the expression data into a finite number of states. Discretization provides clear and interpretable information about the miRNA behaviour and makes the learning process from the data more efficient [7]. However, the procedure is also related to a potential information loss. We comment on this issue in Section 5.

We define a data set *D* = (*S, A*) as a finite set of samples *S* ⊆{0, 1}^*m*^, where *m* ∈ ℕ is the number of miRNAs and *A*: *S* → {0, 1} is sample annotation. The first column includes unique sample IDs and the second the annotation of samples, where 0 is assigned to negative class samples and 1 to positive. The following columns are miRNA expression profiles that describe the miRNA regulation among the samples. miRNAs are binarized into two states: up-regulated (1/positive) and down-regulated (0/negative), according to a given threshold. An example of a data set is presented in Table 1.

**Table 1:** The data: miRNA expression profiles

| ID | Annots | *miR-a* | *miR-b* | *miR-c* |
| --- | --- | --- | --- | --- |
| 1 | 0 | 0 | 1 | 0 |
| 2 | 0 | 0 | 1 | 0 |
| 3 | 1 | 1 | 0 | 0 |
| 4 | 1 | 0 | 0 | 0 |
| 5 | 1 | 1 | 0 | 0 |

A miRNA is non-regulated when for every sample its state is either 0 or 1 (e.g., Table 1, *miR*-*c*). Some miRNAs can perfectly separate the samples into the two categories implied by the annotation (e.g., Table 1, *miR*-*b*). In the following section we introduce single-circuit classifiers that process miRNAs as inputs to classify cell states.

### 2.2 Single-circuit Classifier

A single-circuit cell classifier may be represented by a boolean function *f*: *S* → {0,1}. To make a classifier feasible to construct in the lab additional constraints must be imposed on the function. We adopt here the constraints introduced by Mohammadi et al. [16]. Accordingly, the function should be given in *Conjunctive Normal Form* (CNF), i.e., a conjunction of clauses where each clause is a disjunction of negated (negative) or non-negated (positive) literals. It may consist of: (i) negative literals only in 1-element clauses (NOT gates) (ii) at most 3 positive literals per clause (OR gate) (iii) up to 10 literals and up to 6 clauses in total (iv) including at most 4 NOT gates and 2 OR gates. An example of a classifier satisfying the above-mentioned constraints is presented below.

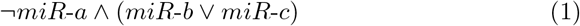

The function should output 1/True in case of cancerous and 0/False in case of healthy cells. The example function presented in Eq. 1 classifies a cell as positive/1 if *miR*-*a* is down-regulated and at least one of the other miRNAs (*miR*-*b* or *miR*-*c*) is up-regulated.

### 2.3 Distributed Classifier

Here, we introduce a concept of *Distributed Classifier* (*DC*) for the cell classification problem. A DC is a finite set *DC* = {*f*_1_, …, *f*_*c*_}, where *f*_*i*_ is a boolean function *f*_*i*_: *S* → {0,1}, to which we will refer from now on as a *Rule*, *c* ≤ *c*_*max*_, *c* ∈ ℕ is the *DC* size and *c*_*max*_ ∈ ℕ is an upper bound for the DC size. Motivated by Section 2.2 a *Rule* must be a boolean function in a *Conjunctive Normal Form* consisting of at most two single-literal clauses. An example of a *DC* is presented below.

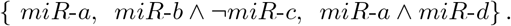

We assume that each *Rule* in the set must be unique, i.e., we do now allow copies of *Rules* in the *DC*. Also, two identical miRNA IDs cannot occur in one *Rule*. The DC categorizes cells according to a threshold function *F*_*DC*_: *S* → {0, 1} with

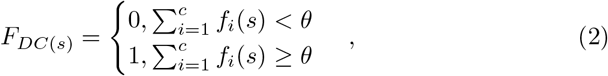

where *s* ∈ *S* is a sample and *θ* ∈ [0, *c*] is a threshold. Here, we use *θ* = ⌊*α* ⋅ *c*⌉ as the threshold, where *α* is a ratio that allows to calculate the decision threshold based on the classifier size. The threshold is then rounded half up. *F*_*DC*_ returns 1/True if a certain number of *Rules* (*θ*) outputs 1/True, e.g., *α* = 0.5 for *c* =5 indicates that at least 3 *Rules* must output 1/True to classify a cell as positive.

Depending on *α* one may receive different results. In case of a very low threshold, e.g., if only one *Rule* outputing 1/True results in *DC* outputing 1/True, the DC becomes simply a disjunction of *Rules*. Note, that the function may then classify in favor of the positive class, as the decision to classify a sample as positive is in fact made by only one rule. This effect is already reduced by not considering 2-literal OR gates as rules. Otherwise, if the threshold is very high, i.e., all the rules must output 1 for the DC to output 1, the function takes a form of a conjunction of clauses staying close to the single-circuit classifier. Unlike the disjunction, a conjunction may classify in favor of the negative class which may decrease the sensitivity of the method. Applying intermediate thresholds results in different combinations of those functions, therefore, different classification performance. In terms of cell classifiers applied as a cancer treatment, one may consider a following problem: in case of high *α*, the classifier may misclassify the diseased cells resulting in false negatives. Thus, the treatment may be less effective. However, low *α* may result in misclassification of healthy cells which makes the treatment more toxic as the drug is released in those cells (false positives). Here, one should consider what type of errors is less desirable and apply a suitable threshold. We discuss this issue further in Section 4.

### 2.4 Evaluation

Here, we introduce the measures we employ to evaluate DCs in terms of their binary classification. Many metrics that may be applied are available [21]. However, real-world expression data is often heavily imbalanced, i.e., the samples are not equally represented in the two classes. Data imbalance may significantly influence the classification results [28]. Balanced Accuracy (BACC) is an intuitive and easily interpretable metric that allows to balance the importance of samples in both classes (Eq. 3) [21]. Thus, as a main measure of classifier’s performance we apply BACC.

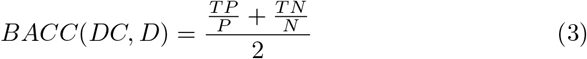

To evaluate other aspects of classification correctness we employ additional common metrics such as sensitivity (*TP/*(*TP* + *FN*)), specificity (*TN/*(*TN* + *FP*)) and accuracy ((*TP* + *TN*)*/*(*T* + *N*)). Sensitivity represents the ability of the method to correctly distinguish samples belonging to the positive class, while specificity represents those belonging to the negative class. Accuracy gives information about the proximity of results to the true values, but does not take data imbalance into account.

## 3 Genetic Algorithm

In this section we present the architecture of a GA applied to design and optimization of DCs. In the following sections we describe the core structure of the algorithm (Algorithm 1) as well as the used parameters and operators. Detailed algorithms, related to, e.g., particular operators, may be found in the Appendix.

### 3.1 General description

The input miRNA expression data must be formatted as described in Section 2.1. To optimize the DCs, seven parameters must be specified: *iter* - number of iterations, *ps* - population size, *cp* - crossover probability, *mp* - mutation probability, *ts* - tournament size, *c*_*max*_ - maximal size of a classifier, *α* - the decision threshold ratio. As output, the algorithm returns a list of all best solutions found over the GA’s iterations according to their balanced accuracy (*DC*_*best*_). In case of single-circuit classifiers, besides the accuracy, the complexity of a solution is also taken into account [16, 1]. Thus, we choose the shortest solution (consisting of the lowest number of rules) as the optimal one.

#### Algorithm 1: A genetic algorithm for *DC* design

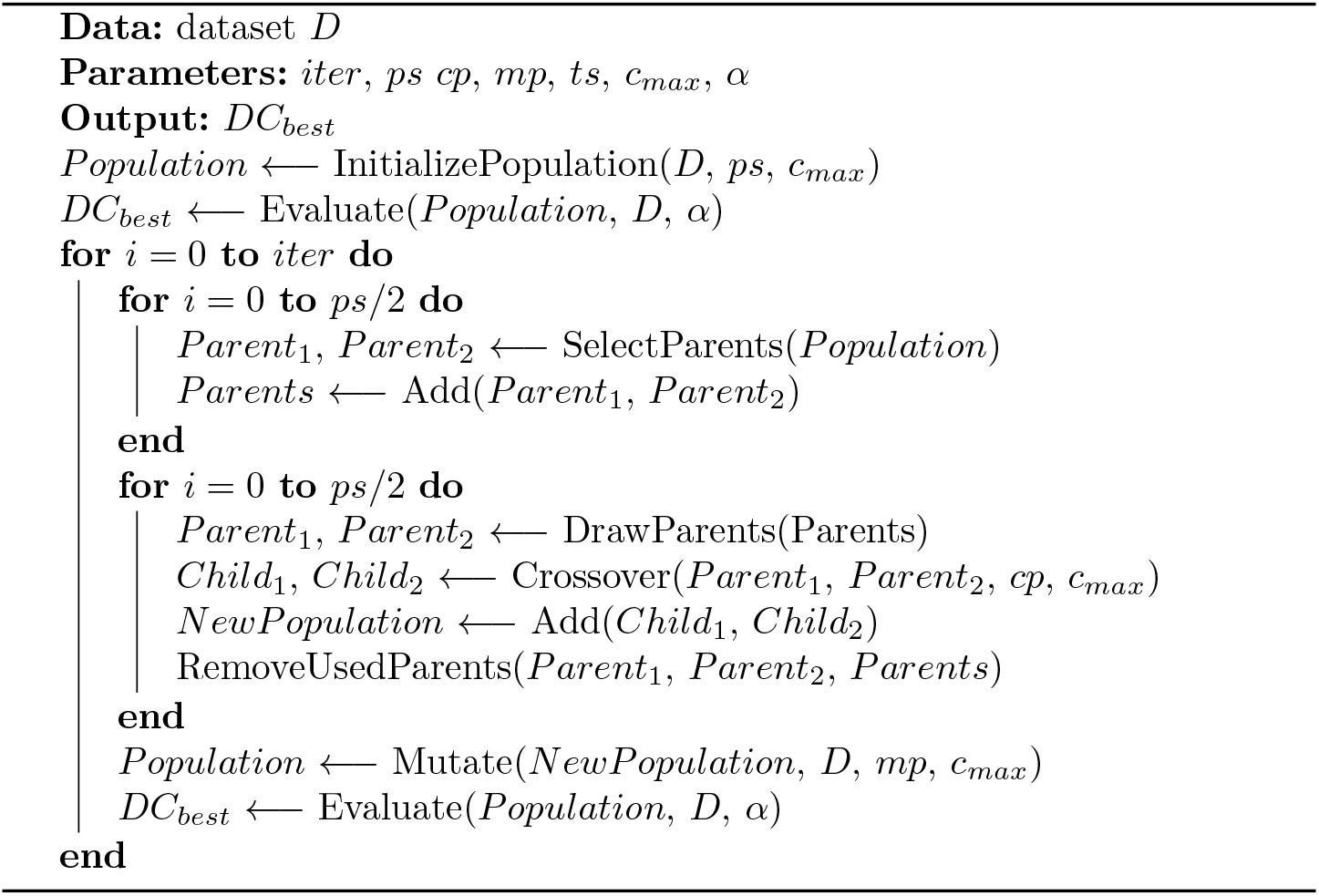

The algorithm starts with a random generation of an initial population. Next, the population is evaluated and a list of best solutions *DC*_*best*_ is created. Having the initial population generated, the algorithm starts with a first generation. At the beginning, *ps* individuals are selected as potential parents to be recombined. Next, the crossover occurs with the probability *cp*. As classifier sizes may differ, we propose two recombination strategies described further in section 3.4. Next, individuals in the new population may mutate with the probability *mp*. The population is then evaluated, i.e., list *DC*_*best*_ of best solutions is updated. All the described steps in a generation are repeated *iter* times. Below we explain the details of the algorithm design.

### 3.2 Fitness Function and Evaluation

As described in Section 2.4, to evaluate the classification performance of a distributed classifier we apply balanced accuracy as the fitness function. To count TPs and TNs we iterate over samples and evaluate the performance of a classifier according to the threshold function described in Section 2.3. The classification threshold is specified by the user. In Section 4 we discuss the influence of different thresholds on the results.

### 3.3 Population

#### Individual Encoding

An individual (i.e., a DC) is encoded as a vector of single rules. A unique ID and a fitness score is assigned to each individual. Both, the distributed classifier and single rules must satisfy previously described constraints (see Section 2.3). Note, rules must consist of unique miRNA-inputs and DCs must consist of unique rules.

#### Initial Population

An initial population of a given size (*ps*) is generated randomly, i.e., each classifier and each single rule in the classifier is randomly initialized. Individuals in the population may be of a different size *c* (maximally *c*_*max*_). Thus, to generate a new individual, *c* must be first defined. Then, each single rule is generated in a few steps. First, the rule size is chosen. Next, numbers of positive and negative inputs in a rule are defined. Finally, miRNA IDs are assigned to the inputs. For more details see Appendix (Algorithm 2).

### 3.4 Operators

#### Selection

Parents, to be potential candidates for recombination, are chosen in a process of *tournament selection*. Many selection operators are described in the literature. Tournament selection allows to maintain diversity in the population and can be efficiently implemented [24]. In each selection iteration two parents are chosen in separate tournaments. To select one parent, a number of *ts* individuals is randomly chosen from the current population to participate in a tournament. The winning candidate is an individual with the best fitness score. The first and the second parent must be different individuals. Thus, in each iteration, after choosing the first parent, its ID is temporarily blocked to be re-selected. The steps are repeated to form a population of selected individuals of the size of the original population (*ps*). For more details see Appendix (Algorithm 3).

#### Crossover

In each crossover iteration two parents are randomly chosen from a population of selected individuals to recombine. Crossover occurs with the probability *cp*. To decide whether parents exchange information a random number *p* is chosen. If *p ≤ cp* then the parents recombine to generate offspring. Otherwise, parents are copied to a new population. As individuals may have different sizes, we apply two strategies. If chosen parents are of the same size we perform standard uniform crossover. Otherwise, to preserve equal chance for each rule to be swapped, we apply index-based crossover. The *crossover index* is chosen randomly and allows to change the position of a shorter individual in relation to the second one. Here, the rules may be either swapped or, if there is no possibility to swap them due to different sizes, a rule from a larger classifier may be copied to the offspring. Note, that the index-based crossover may shorten the size of an individual as additional rules cannot be added to the larger classifier. After the crossover, the parents are removed from the population of selected individuals. For more details see Appendix (Algorithm 4 and 5).

#### Mutation

Mutation may occur on two levels: both, rules and inputs may mutate. A rule may (i) be removed from a classifier, (ii) be added to a classifier and (iii) be copied from one classifier to another. As mentioned before, index-based crossover may shorten the classifier. Here, two possibilities to extend the size of a classifier are available: a new rule may be initialized and added to a classifier or copied from another classifier. These two options balance the influence of crossover on the size of classifiers. An input may (i) be removed from a rule, (ii) be added to a rule, (iii) may change the sign (i.e., become a negative or positive input respecting the constraints described in Section 2.3). Rules, being larger components affecting the classifier size, mutate with a lower probability than inputs (0.2). Note, the maximal size of a classifier (*c*_*max*_) must be preserved. For more details see Appendix (Algorithm 6).

## 4 Case Studies

In this section, we illustrate the potential of DCs in application, in particular, in cancer cell classification by performing case studies on real-world breast cancer data. We first describe the data sets used to evaluate DC performance. Then, we present results of parameter tuning and cross-validation. We analyze classifier performance, as well as the relevance of chosen miRNAs. Finally, we compare DCs with a single-circuit classifier design approach.

### 4.1 Breast Cancer Data

To evaluate the performance of our approach we use Breast Cancer data sets previously applied by Becker et al. [1] and Mohammadi et al. [16] to the design of single-circuit classifiers. Originally the data was described by Farazi et al. [6] and pre-processed by Mohammadi et al. [16]. The details about the samples and miRNAs may be found in Table 2. The data set All includes samples of different breast cancer subtypes. This allows to compare breast cancer samples with the control (negative samples). The following data sets are subsets representing different breast cancer subtypes containing information about the differences between particular subtypes and the control. Note, the data sets are significantly imbalanced as the negative class is heavily underrepresented. The data is formatted according to the description presented in Section 2.1. In terms of cell classifiers, non-regulated miRNAs do not carry any information. Thus, we remove them from the data sets before optimizing the classifiers. The last two columns of Table 2 include numbers of miRNAs before and after the filtering procedure.

**Table 2:** Breast Cancer data description.

| Dataset | Samples | Positive | Negative | miRNAs | filtered miRNAs |
| --- | --- | --- | --- | --- | --- |
| All | 178 | 167 | 11 | 478 | 57 |
| Triple- | 82 | 71 | 11 | 456 | 52 |
| Her2+ | 86 | 75 | 11 | 438 | 19 |
| ER+ Her- | 32 | 21 | 11 | 392 | 18 |
| Cell Line | 17 | 6 | 11 | 375 | 59 |

### 4.2 Parameter tuning

To tune the parameters of the genetic algorithm we applied a random search approach. Random search methods allow to obtain results similar to the grid search approach, while decreasing the computational cost [2]. This provides an opportunity to extend the range of tested parameters. To tune the parameters we used the Breast Cancer All data set. We performed 3-fold cross-validation and repeated each GA run 10 times to obtain the average balanced accuracy on the test data. We have randomly chosen 300 combinations of 5 parameters in following ranges: *iter*: 25 −100, step 25; *ps*: 50 - 300, step 50; *cp*: 0.1 - 1.0, step 0.1; *mp*: 0.1 - 1.0, step 0.1; *ts*: 10 - 50%, step 10% (of *ps*). We tuned the parameters for *α* = 0.50 and *c*_*max*_ =5 and chose a following set of parameters based on average scores: *iter* = 75, *ps* = 200, *cp* = 1.0, *mp* = 0.3, *ts* = 10% (20 individuals). We apply those parameters to all case studies presented in the following sections.

### 4.3 Cross-validation

To evaluate the classifiers accuracy we performed 3-fold cross-validation for the breast cancer data sets presented in Section 4.1. We divided the data sets in 3 folds nearly equal in terms of the number of samples representing each class per fold. For all tests we apply *c*_*max*_=5. The classifier size *c*_*max*_=5 allows to preserve the maximal number of miRNA inputs as proposed for single-circuit classifiers [16, 1]. Maintaining similar complexity of classifiers allows to compare the DC-based method to another approach.

We test eight different values of *α*: 0.25, 0.35, 0.40, 0.50, 0.60, 0.65, 0.75, 0.85, to evaluate the influence of the threshold function on the classification accuracy. The best results are presented in Table 3 (complete results for different *α* values may be found in the Appendix, Table A2). The DCs presented in the results are the first best shortest classifiers found by the algorithm. If identical BACC values for the testing data were obtained for more than one *α*, we present results for a DC with the highest BACC value on the training data. Otherwise (equal training BACC values), we present an exemplary result for a chosen threshold. Table 3 includes the *α*-s and performance scores. All scores except of BACC (train) were calculated on the testing data.

**Table 3:** Results of 3-fold cross-validation. For the Breast Cancer All data set we found DCs performing with identical score values for two *α* values (0.50, 0.60) and for ER+Her- for 6 different *α* values (0.35, 0.50, 0.60, 0.65, 0.75, 0.85)

| Dataset | $\alpha$ | Sensitivity | Specificity | ACC | BACC | BACC<sub>train</sub> |
| --- | --- | --- | --- | --- | --- | --- |
| All | 0.50 | 0.92 | 0.92 | 0.92 | 0.92 | 0.98 |
| Triple- | 0.85 | 0.92 | 0.75 | 0.89 | 0.83 | 0.98 |
| Her2+ | 0.75 | 0.99 | 0.61 | 0.94 | 0.80 | 0.96 |
| ER+ Her- | 0.50 | 0.90 | 0.64 | 0.82 | 0.77 | 0.93 |
| Cell Line | 0.25 | 1.00 | 1.00 | 1.00 | 1.00 | 1.00 |

High BACC values obtained for the training data sets, as well as the average final population BACC values (0.91), show that the populations converge over the iterations resulting with high-performing DCs. The BACC values measured for the testing data sets are significantly higher for the largest and the smallest data sets than for the intermediate-size ones. The accuracy is higher than BACC for all data sets as the metric is not sensitive to data imbalance.

The sensitivity is high for all data sets meaning that the method successfully classifies samples belonging to a positive class. However, the specificity is decreased for intermediate-size data sets. Note, the data sets are significantly imbalanced, i.e., the negative class is strongly underrepresented. Thus, even small number of errors results in substantially decreased specificity.

**Table 4:** Comparison of results of 3-fold cross-validation for the ASP-based approach proposed by Becker et al. [1] and for the GA for *α* values presented in Table 3.

| Dataset | Method | Sensitivity | Specificity | ACC | BACC | BACC<sub>train</sub> |
| --- | --- | --- | --- | --- | --- | --- |
| All | GA | 0.92 | 0.92 | 0.92 | **0.92** | 0.98 |
|  | ASP | 0.96 | 0.47 | 0.93 | 0.72 | 0.92 |
| Triple- | GA | 0.92 | 0.75 | 0.89 | **0.83** | 0.98 |
|  | ASP | 0.89 | 0.44 | 0.83 | 0.67 | 0.96 |
| Her2+ | GA | 0.99 | 0.61 | 0.94 | 0.80 | 0.96 |
|  | ASP | 1.00 | 0.61 | 0.95 | **0.81** | 0.96 |
| ER+ Her- | GA | 0.90 | 0.64 | 0.82 | **0.77** | 0.93 |
|  | ASP | 0.90 | 0.64 | 0.82 | **0.77** | 0.93 |
| Cell Line | GA | 1.00 | 1.00 | 1.00 | **1.00** | 1.00 |
|  | ASP | 0.83 | 1.00 | 0.93 | 0.92 | 1.00 |

The best *α* values differ among the data sets. For the largest one, *α* is equal or not much higher than 50%. The data sets of intermediate sizes (Triple- and Her2+) favoured two more extreme *α* values. For the ER+Her-several *α* values returned identical results (Appendix, Table A2). For the smallest data set the lowest *α* value resulted in the highest BACC. Thus, the threshold seems to be data-specific and should be adjusted to the data set for the DC to perform well.

Applying a certain threshold caused a shift in average numbers of certain types of errors. Here, we analyze average numbers of FPs and FNs observed among all data sets for two extreme applied *α* values. As expected, in case of a high threshold (0.85) the shift is displayed towards misclassification of the positive samples (*FP*_*avg*_ = 1.00, *FN*_*avg*_=2.00). Otherwise, the low threshold (0.25) causes more frequent misclassification of negative samples (*FP*_*avg*_ = 1.20, *FN*_*avg*_=0.87). Thus, one may take it into account while designing a classifier. Complete information about average FPs and FNs for different thresholds may be found in the Appendix, Table A1.

All the tests were performed using Allegro CPU Cluster provided by Freie Universitaet Berlin^3^. An average runtime for one fold of cross-validation for all the data sets employed in the case studies is 25 min 42 sec. Thus, the tests may be also performed on a personal computer. However, the breast cancer data sets consist of up to 180 samples and about 50 relevant miRNAs. Thus, one should consider performing extended scalability tests to estimate the runtime limits of the method.

### 4.4 Analysis of input viability

In this section we analyze miRNA inputs that occur in two exemplary classifiers. We chose the best performing classifiers for the largest data set (All) representing all subtypes and the smallest Cell Line data set.

For breast cancer All two different *α* values resulted in the highest BACC. We found that classifiers for each cross-validation fold in the data set are identical for both *α* values. Also, all the classifiers are of the same size *c* = 4. In this case the applied *α* does not change the threshold function between both values (0.50 and 0.60), i.e., for all data sets at least 2 *Rules* must output 1 to classify a cell as positive. Below we present a DC found for the third cross-validation fold of the All data set. The classifier consists of 4 different 1-input rules. We analyzed the miRNAs and found that all of them may be relevant for cancer sample classification. The classifier is presented below.

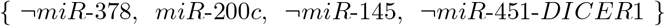

*miR*-378, *miR*-145, *miR*-451-*DICER*1 are described as down-regulated in breast cancer [4, 6], e.g., the study by Ding et al. [4] has shown that underexpression of *miR*-145 is related to increased proliferation of breast cancer cells. Also, miR-378 occurred as down-regulated in the best 1-input single-circuit classifier presented previously by Becker et al. for the same data set [1]. miR-200c is marked as up-regulated in breast cancer in [22].

Another classifier we present is a DC for the third cross-validation fold for the Cell Line data set:

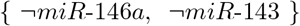

For most of the *α* values the performance of found *DC*s was very low for this particular fold in the Cell Line data set (BACC = 0.50). A perfect classifier of size 2 performing with BACC = 1.00 on both training and testing data was found with *α* = 0.25, i.e., one of 2 rules must output 1 to classify the cell as positive. We found that both, miR-146a and miR-143, are described as down-regulated in breast cancer [11, 17].

### 4.5 Comparison to other methods

We optimized single-circuit classifiers with the ASP-based method proposed by Becker et al. [1] by performing 3-fold cross-validation using the same data sets and identical division into folds. The objective function of the ASP algorithm is based on the minimization of the total number of classification errors. Note that the ASP method may return several optimal classifiers. Different combinations of FPs and FNs influence Balanced Accuracy. Thus, to increase the chance of ASP to perform well, we have chosen the best classifiers according to their BACC. Here, we do not compare our results to Mohammadi et al. [16] as the approach did not perform better than the ASP-based approach as described by Becker et al. in terms of binary classification [1].

The DC-based method outperformed the single-circuit approach in 3 of 5 case studies. For two other data sets the resulting BACC (test) values are either identical (ER+Her-) or very similar (Her2+). This may imply that further improvement of classifier performance for those data sets is not possible with the currently applied techniques. The training BACC values are also significantly higher for the DC-based approach. Note, the DC-based design method explores a different search space than the single circuit approach. Although single circuits are also allowed as 1-rule classifiers, their complexity is substantially lower in comparison to single circuits. Additionally, ASP returns globally optimal solutions, i.e., it adjusts the classifier perfectly to the training data, which may cause overfitting. Although, the classifiers obtain high BACC on the training data (average for all data sets: 0.95), the classifiers may be too specific to perform well on the testing data.

## 5 Discussion

In this article, we introduced a new approach to cell classifier design. The concept of DCs proposed by Didovyk et al. [3] was re-formalized in the context of miRNA-based cell classification. We designed and implemented a genetic algorithm that allows design and optimization of *DC*s. We performed case studies on real-world data and compared our results to a single-circuit design method obtaining significantly higher or similar accuracy.

DCs show immense potential as an alternative to single-circuit designs. Presented case studies demonstrate the DC’s ability to perform classification on real-world cancer data. The results obtained on the training data show that the proposed genetic algorithm allows to optimize classifiers that achieve high accuracy. The cross-validation demonstrates that the optimized DCs classify unknown data with high accuracy. Although, the algorithm performs better on the largest and the smallest data sets than on the intermediate-size ones, the results for Her2+ and ER+Her-are very similar which may suggest that for those data sets significant improvement is not possible. The data sets for which the algorithm returns the worst results (Her2+, ER+Her-) are ones with the lowest number of relevant miRNAs. Thus, the number of possible solutions is significantly decreased in contrast to other data sets. The comparison to a single-circuit design method shows that DCs outperformed single-circuit classifiers on most of the presented data sets according to balanced accuracy. The improvements in binary classification may be a result of applying a different strategy to cell classifier design. Here, single-circuit decision is complemented by a collective classification based on a threshold function. Thus, the DCs may be more resistant to data noise than single-circuit classifiers.

Generally, the problem of designing reliable and efficient DCs begins with the initial data processing. As mentioned before, the data sets employed for the case studies are significantly imbalanced. Although we apply an objective function that allows to partially overcome this issue, one may consider applying data balancing methods such as weighted schemes that balance the sample importance [8].

Our approach to the design of DCs is based on binarized data sets. As mentioned before, data discretization allows obtaining clear-cut information about miRNA regulation and efficient exploration of the search space. One advantage of this data processing procedure is absorption of noise coming from, e.g., lab artifacts. However, simultaneously some information that may be valuable for the classification is lost. Considering binarization according to a given threshold, miRNAs having their concentrations significantly higher (or lower, respectively) than the threshold may be more informative. Thus, one may introduce a multi-objective function that allows to optimize both, the accuracy and the use of particular miRNAs according to, e.g., a weighted scheme favoring more reliable miRNAs.

Adapting the ASP approach to single circuit classifier design, one could apply ASP to the optimization of DCs, obtain globally optimal solutions and compare with the heuristic approach. However, ASP searches through the entire solution space; thus, the runtime may be significantly increased with the rising number of possible combinations. As we expect that this may significantly limit the ASP-based optimization, one may explore other possibilities. In the proposed GA the initial population is generated randomly, i.e., there is no preference in choosing particular miRNAs or gate signs that built rules. One may optimize the initial population by creating rules taking such preferences into account. The ASP allows to optimize single short classifiers with relaxed constraints in a short time, e.g., allowing up to a certain number of errors. This may generate a pool of rules that are pre-optimized resulting in a better starting point for the algorithm.

Although the results demonstrate that classifiers perform with high accuracy, the possibilities to further develop the presented method should be explored. Certainly, the approach must be tested using more data representing variety of cancer types. Although the proposed genetic algorithm performed well on the presented case studies, particular parts of the algorithm may be improved. One may consider to extend the parameter tuning procedure by applying more sophisticated methods such as GAGA approach using a metaGA above the main algorithm to tune the parameters [5]. Also, different selection operators may be tested to evaluate the influence of a chosen operator on the results [24]. Although tournament selection is described in the literature as a well-performing operator, some other operators may be more accurate for particular problems than the commonly recommended ones.

Although DCs are not yet applied in terms of cancer cell classification, the approach should be further investigated. DCs are designed based on available building blocks that are in fact single-circuit classifiers. Mohammadi et al. [16] presented a biochemical model of a single-circuit classifier that allows to manipulate the output compound concentration. Thus, the biological output threshold for a given classifier may be adjusted to perform the classification in living cells. As the on-off single-circuit response may be regulated on the biological level, the sum of their outputs should also be adaptable for a given DC. This needs to be investigated through further work in the lab.

## Data and Software Availability

The algorithm is implemented in Python 3. The scripts, as well as the data used to tune the parameters and test the algorithm’s performance including the results, are available at GitHub [19].

## Aknowledgements

We would like to thank to Pejman Mohammadi, Yakoov Benenson and Niko Beerenwinkel for sharing the breast cancer data with us. MN would like to thank to Jakub Bartoszewicz (Robert Koch Institute, Berlin) for his valuable comments and support with cluster handling.

## Appendix

### Algorithm 2: Population initialization

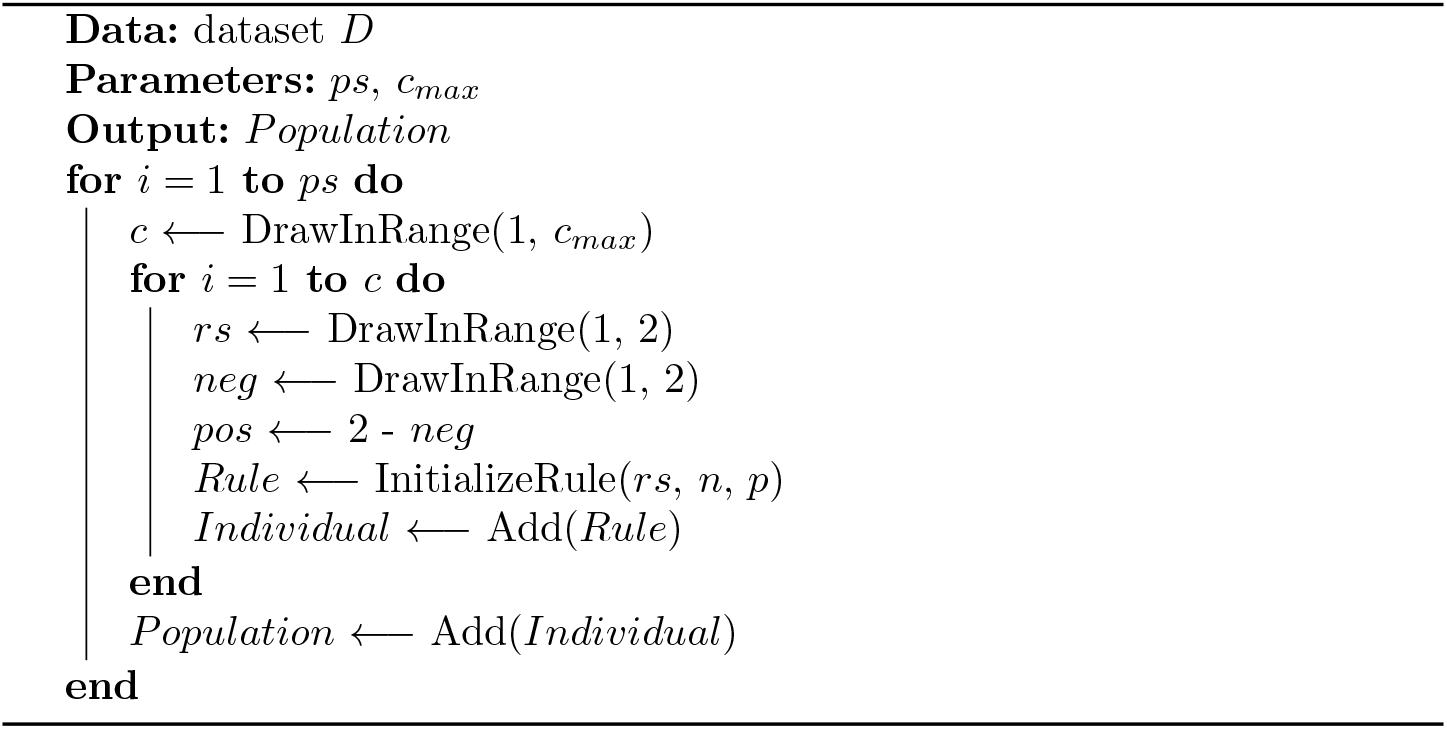

### Algorithm 3: Selection of parents

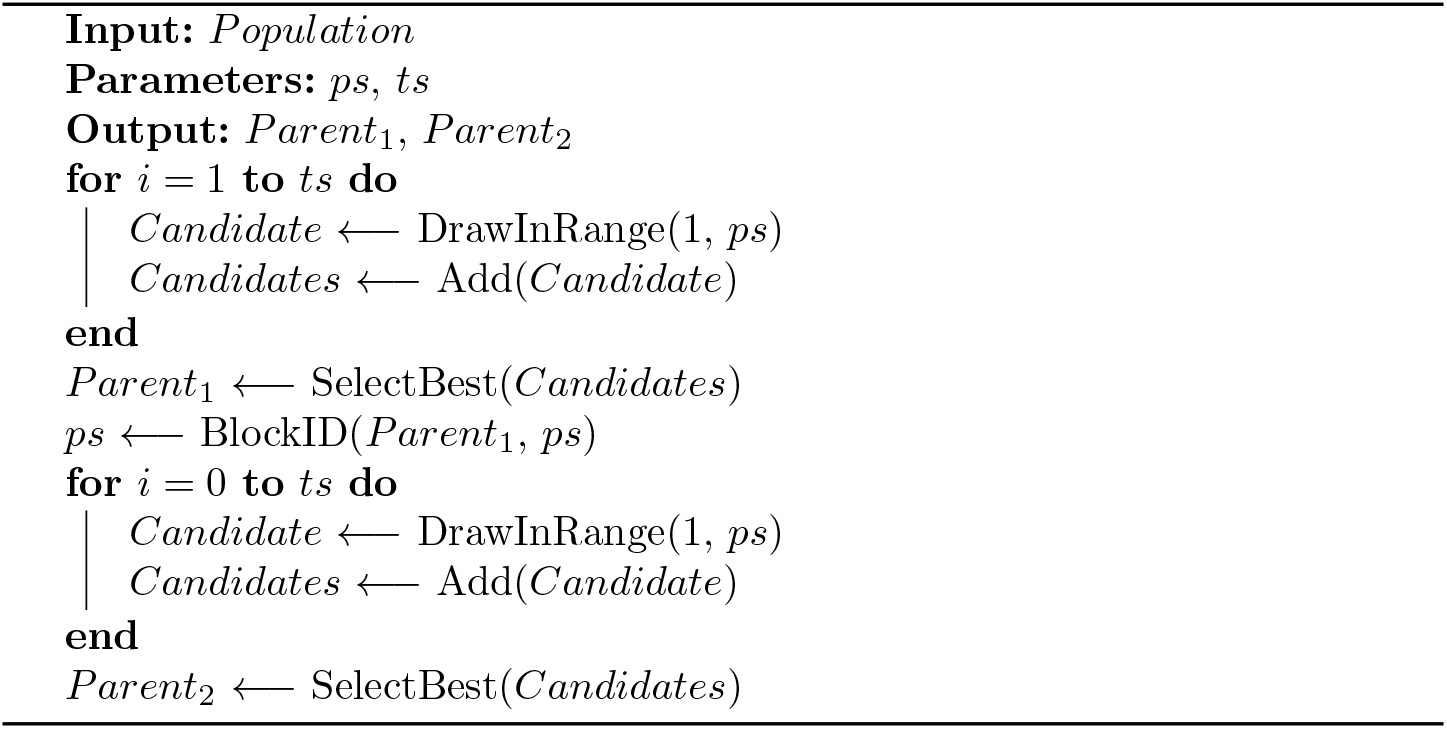

### Algorithm 4: Crossover

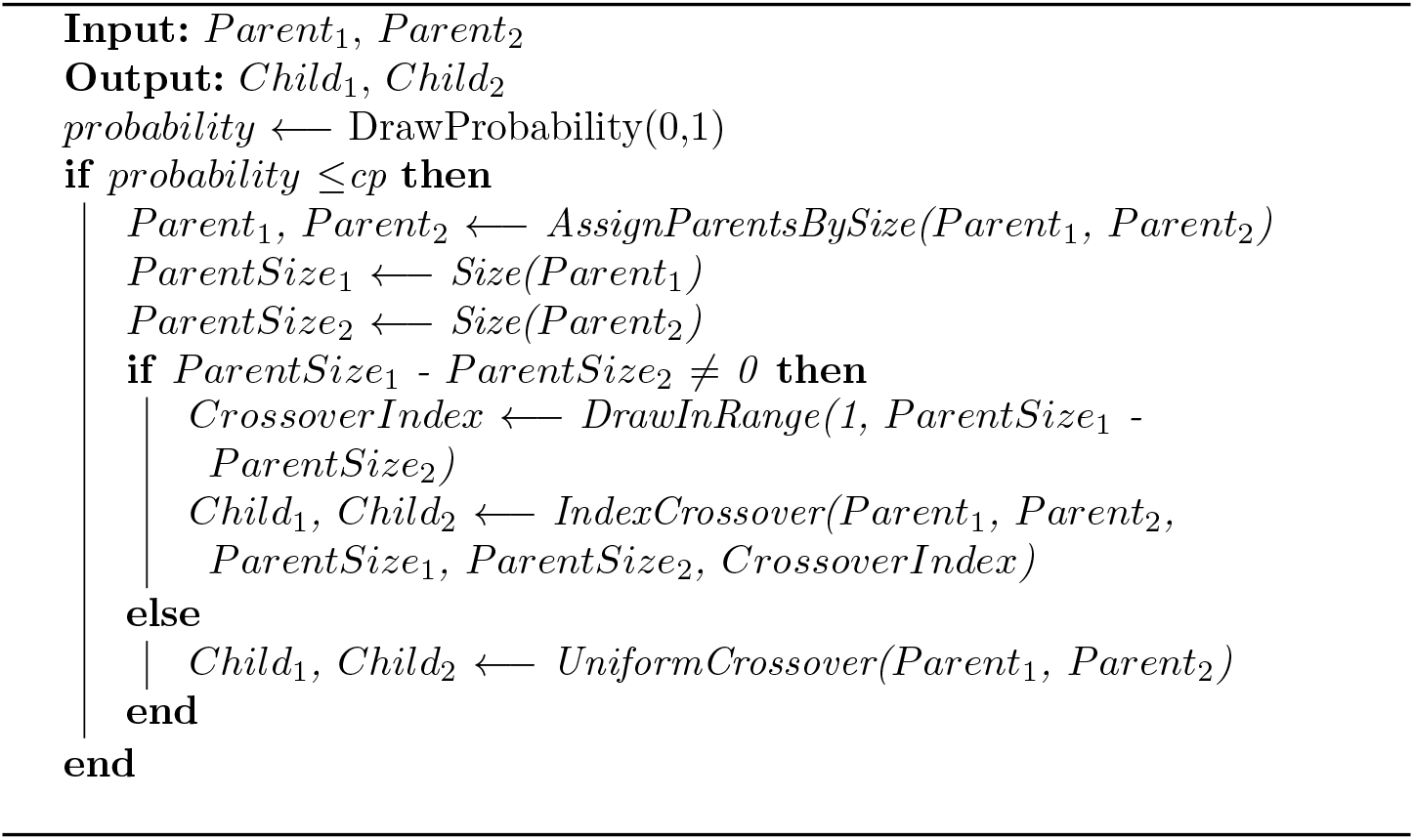

### Algorithm 5: Index-based crossover

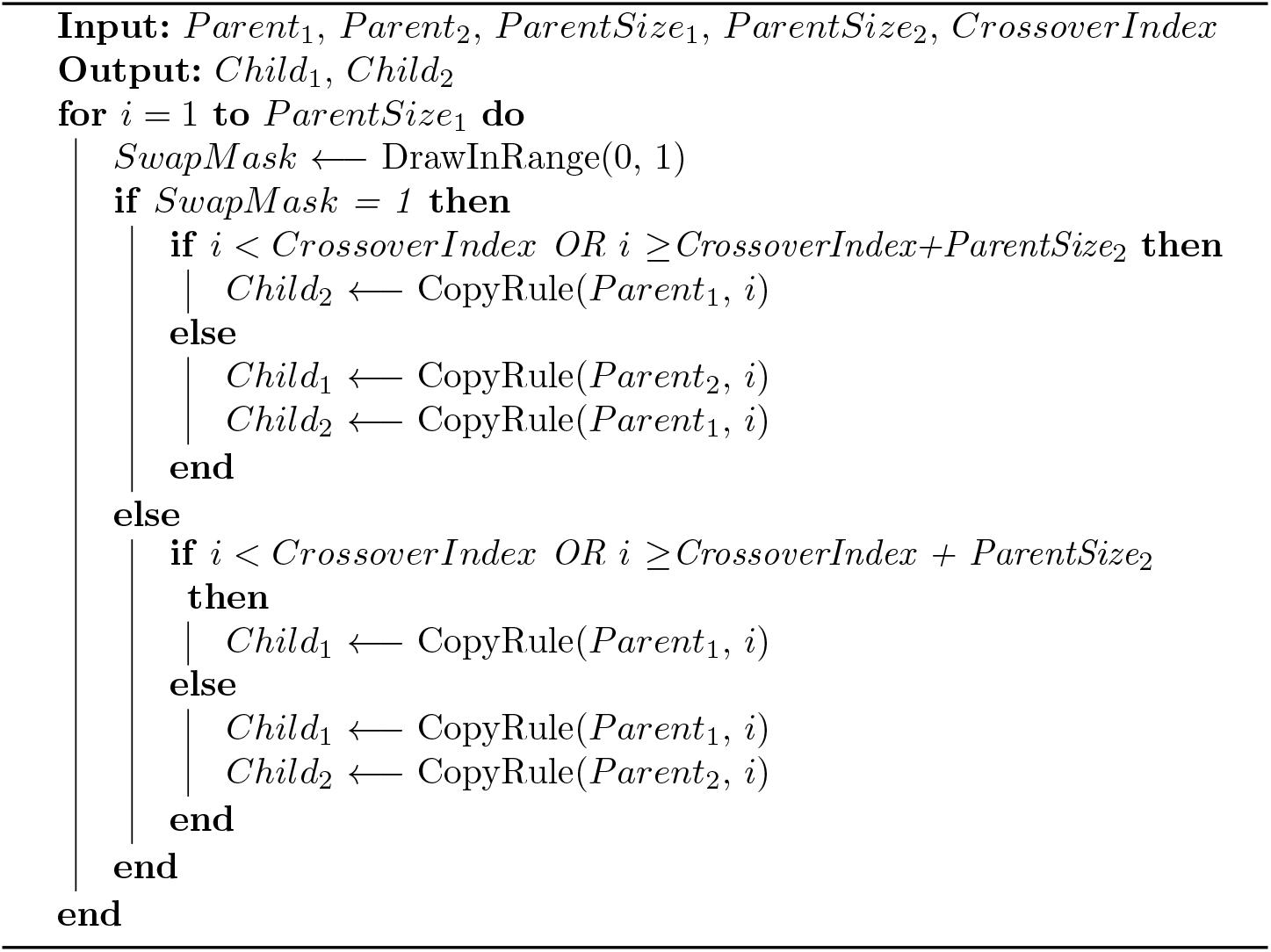

### Algorithm 6: Mutation

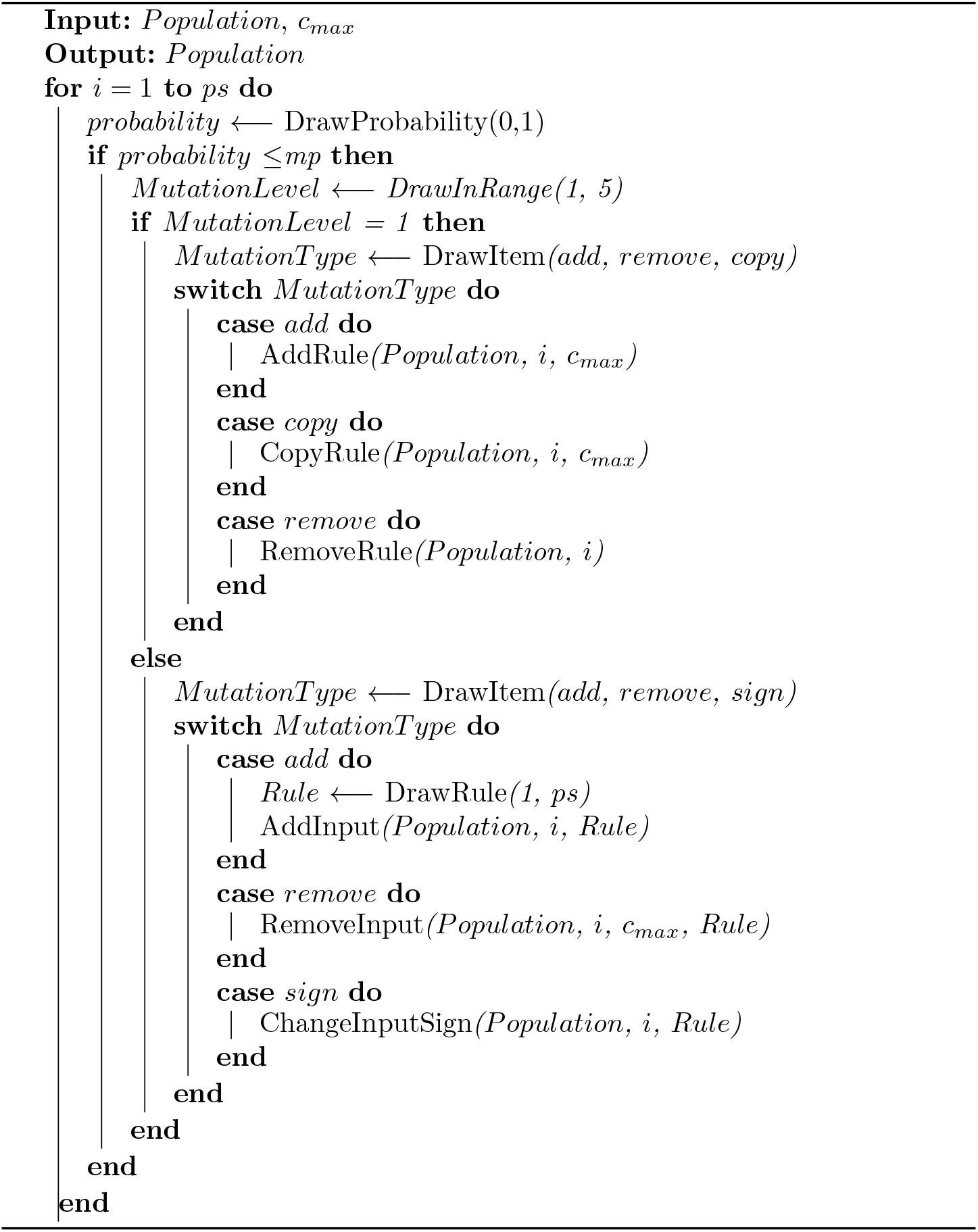

**Table A1:** Average number of FPs and FNs for different thresholds (for all datasets).

| Threshold | $FP_{avg}$ | $FN_{avg}$ |
| --- | --- | --- |
| 0.85 | 1.00 | **2.00** |
| 0.75 | 0.93 | **1.20** |
| 0.65 | 1.13 | **1.20** |
| 0.60 | 0.93 | **1.47** |
| 0.50 | 0.93 | **1.47** |
| 0.40 | **1.27** | **1.27** |
| 0.35 | **1.20** | 0.73 |
| 0.25 | **1.20** | 0.87 |

**Table A2:** Results of 3-fold cross-validation.

| Dataset | $\alpha$ | Sensitivity | Specificity | ACC | BACC | BACC<sub>train</sub> |
| --- | --- | --- | --- | --- | --- | --- |
| All | 0.85 | 0.89 | 0.83 | 0.88 | 0.86 | 0.96 |
|  | 0.75 | 0.94 | 0.81 | 0.93 | 0.87 | 0.98 |
|  | 0.65 | 0.95 | 0.72 | 0.93 | 0.83 | 0.98 |
|  | 0.60 | 0.92 | 0.92 | 0.92 | **0.92** | 0.98 |
|  | 0.50 | 0.92 | 0.92 | 0.92 | **0.92** | 0.98 |
|  | 0.40 | 0.94 | 0.64 | 0.92 | 0.79 | 0.99 |
|  | 0.35 | 0.97 | 0.72 | 0.96 | 0.85 | 0.99 |
|  | 0.25 | 0.96 | 0.72 | 0.94 | 0.84 | 1.00 |
| Triple- | 0.85 | 0.92 | 0.75 | 0.89 | **0.83** | 0.98 |
|  | 0.75 | 0.96 | 0.67 | 0.92 | 0.81 | 0.99 |
|  | 0.65 | 0.94 | 0.58 | 0.89 | 0.76 | 1.00 |
|  | 0.60 | 0.93 | 0.64 | 0.89 | 0.78 | 1.00 |
|  | 0.50 | 0.93 | 0.64 | 0.89 | 0.78 | 1.00 |
|  | 0.40 | 0.94 | 0.56 | 0.89 | 0.75 | 1.00 |
|  | 0.35 | 0.94 | 0.53 | 0.89 | 0.74 | 0.99 |
|  | 0.25 | 0.94 | 0.53 | 0.89 | 0.74 | 1.00 |
| Her2+ | 0.85 | 0.99 | 0.44 | 0.92 | 0.72 | 0.96 |
|  | 0.75 | 0.99 | 0.61 | 0.94 | **0.80** | 0.96 |
|  | 0.65 | 0.99 | 0.53 | 0.93 | 0.76 | 0.96 |
|  | 0.60 | 1.00 | 0.53 | 0.94 | 0.76 | 0.96 |
|  | 0.50 | 1.00 | 0.53 | 0.94 | 0.76 | 0.96 |
|  | 0.40 | 0.99 | 0.53 | 0.93 | 0.76 | 0.96 |
|  | 0.35 | 1.00 | 0.53 | 0.94 | 0.76 | 0.96 |
|  | 0.25 | 1.00 | 0.53 | 0.94 | 0.76 | 0.93 |
| ER+ Her- | 0.85 | 0.90 | 0.64 | 0.82 | **0.77** | 0.93 |
|  | 0.75 | 0.90 | 0.64 | 0.82 | **0.77** | 0.93 |
|  | 0.65 | 0.90 | 0.64 | 0.82 | **0.77** | 0.93 |
|  | 0.60 | 0.90 | 0.64 | 0.82 | **0.77** | 0.93 |
|  | 0.50 | 0.90 | 0.64 | 0.82 | **0.77** | 0.93 |
|  | 0.40 | 0.90 | 0.53 | 0.78 | 0.72 | 0.93 |
|  | 0.35 | 0.90 | 0.64 | 0.82 | **0.77** | 0.93 |
|  | 0.25 | 0.90 | 0.53 | 0.78 | 0.72 | 0.91 |
| Cell Line | 0.85 | 0.67 | 1.00 | 0.87 | 0.83 | 1.00 |
|  | 0.75 | 0.67 | 1.00 | 0.87 | 0.83 | 1.00 |
|  | 0.65 | 0.67 | 1.00 | 0.87 | 0.83 | 1.00 |
|  | 0.60 | 0.67 | 1.00 | 0.87 | 0.83 | 1.00 |
|  | 0.50 | 0.67 | 1.00 | 0.87 | 0.83 | 1.00 |
|  | 0.40 | 0.67 | 1.00 | 0.87 | 0.83 | 1.00 |
|  | 0.35 | 1.00 | 0.89 | 0.93 | 0.94 | 1.00 |
|  | 0.25 | 1.00 | 1.00 | 1.00 | **1.00** | 1.00 |

## Footnotes

3 https://www.allegro.imp.fu-berlin.de/Cluster

